## Supplementary Information for "Glassy Adhesion Dynamics Govern Transitions Between Sub-Diffusive and Super-Diffusive Cell Migration on Viscoelastic Substrates"

### Supplementary Notes:

#### 1. Dissociation time constant distribution:

The $\tau_{off}$ distribution is sampled from the following power law equation:

$$p(\tau_{off}) = \frac{|(\beta - 1)|}{\tau_{min}} \left(\frac{\tau_{off}}{\tau_{min}}\right)^{-\beta}$$

The parameter β primarily determines the "heaviness" of the distribution's tail, which is a key characteristic of glassy, heterogeneous systems like cell adhesion complexes where broad timescales are essential to capture both frequent short interactions and rare, prolonged interactions. This variance is sensitive to β alone, resulting in higher variability when β<3, which supports the long-tailed distribution that describes the complex temporal dynamics observed in biological adhesion processes.

In contrast, the parameter $\tau_{min}$ sets a lower bound on the dissociation timescale, representing the shortest possible time at which an adhesion bond can dissociate under typical cellular conditions. Physiologically, $\tau_{min}$ corresponds to the smallest timescale needed for initial, quick unbinding events—likely representing transient, weak interactions where bonds are formed but quickly broken under minimal force. By ensuring a minimum timescale, $\tau_{min}$ helps prevent unrealistic, infinitely fast dissociation in the model and reflects the natural constraints imposed by molecular interactions and energy barriers in cell adhesion complexes.

Variance of the distribution is its second moment.

$$\text{Distribution: } p(x) = \frac{\beta}{x_{min}} \cdot \left(\frac{x}{x_{min}}\right)^{-\beta}$$

Second moment:

$$V[x] = \int_{x_{min}}^{\infty} x^2 \cdot \frac{\beta}{x_{min}} \cdot \left(\frac{x}{x_{min}}\right)^{-\beta} dx$$

$$= \frac{\beta}{x_{min}^{1-\beta}} \cdot \int_{x_{min}}^{\infty} x^{2-\beta} dx$$

$$= \frac{\beta}{x_{min}^{1-\beta}} \cdot \left[x^{3-\beta}\right]_{x_{min}}^{\infty}$$

If $\beta \geq 3$, then the second moment converges and is finite.

If $\beta < 3$, then the variance is not finite and do not follow central limit theorem (CLT).

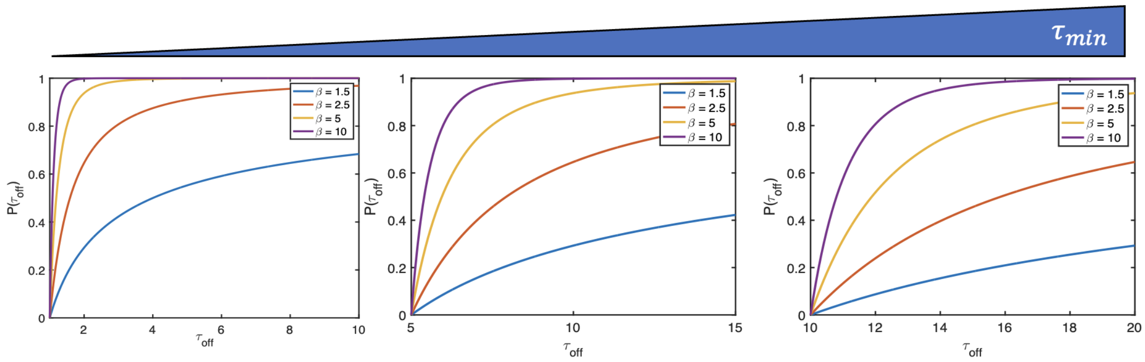


**Supplementary Figure 1:** Effect of long-tailed distribution parameters ($\tau_{min}$ $and$ $\beta$) on the $\tau_{off}$ cumulative distribution.

### 2. Effective off rate constant consisting of unfolding, unbinding and breaking:

In the context of adhesion dynamics and protein mechanics, unbinding, unfolding, and dissociation each represent different processes involving the separation of proteins or conformational changes within them. Here's a breakdown of these terms:

**Unfolding:** Unfolding is the process where a protein loses its specific, folded three-dimensional structure (secondary and tertiary structures) due to applied force or environmental conditions, such as changes in pH or temperature. In adhesion dynamics, unfolding often refers to the stretching and unfolding of proteins like talin and vinculin under force. These proteins can act as molecular clutches: they bind to other proteins, and their unfolding under mechanical load allows cells to maintain traction forces while migrating.

**Unbinding:** Unbinding refers to the process where two proteins or molecular components that are physically interacting separate from each other without structural alteration to either. This process is primarily governed by the strength of the bonds or interactions (like hydrogen bonds, ionic interactions, and van der Waals forces) between the molecules. In focal adhesion complexes, unbinding might describe, for example, the separation of integrins from the extracellular matrix or other associated proteins.

**Dissociation (off-rate):** Dissociation in this context refers to the overall process by which a protein complex or interaction site completely separates, including any sequential steps required for this to occur. For a protein complex that is mechanically loaded, dissociation might involve both unfolding of structural domains (which occurs first) and unbinding of the final remaining contacts between proteins. Dissociation thus represents the culmination of sequential events where force may cause a protein to unfold before ultimately leading to unbinding and complete separation.

In summary, unbinding typically refers to the separation of interacting sites, unfolding describes a conformational change in response to force, and dissociation encompasses the entire sequence, often requiring both unbinding and unfolding in mechanotransducive processes. In adhesion mechanics, capturing the nuances of these events is essential, as each contributes uniquely to cellular responses and migration dynamics. Hence, we can write the effective dissociation rate as:

$$k_{off} = \frac{k_{unbind} * k_{unfold}}{k_{unbind} + k_{unfold}}$$

Now, an effective timescale can be written as:

$$\frac{1}{\tau_{off}} = \frac{\frac{1}{\tau_{unbind} * \tau_{unfold}}}{\frac{1}{\tau_{unbind}} + \frac{1}{\tau_{unfold}}} \tau_{off} = \tau_{unbind} + \tau_{unfold}$$

This relationship allows us to capture the stochastic nature of the unfolding and unbinding processes, which could each independently follow power-law distributions or we can treat unbinding as a constant timescale once proteins are unfolded. The key insight here is that in either case, the effective off-rate constant timescale distribution $\tau_{off}$ inherits power-law characteristics, displaying broad, glassy distributions with heavy tails.

This result is powerful because a power-law timescale distribution naturally aligns with the behavior observed in glassy systems. Specifically, the variance of $\tau_{off}$ can be rendered infinite for power-law exponents $\beta<3$, which promotes truly non-Gaussian, long-tailed distributions capable of capturing diverse adhesion behaviors. Therefore, modeling $\tau_{off}$ as a long-tailed power-law distribution enables us to encapsulate the heterogeneity present in adhesion dynamics, regardless of whether the stochasticity originates predominantly from unfolding or unbinding events. This ensures that our model accurately reflects the complex, glassy nature of adhesion formation and breakage dynamics in cell migration.

**3. Solving the motor clutch model:**
To solve the individual motor clutch module, we adopt the Kinetic Monte Carlo method, which incorporates stochasticity in the cell migration process and is based on out previous work [1]. The detailed steps of the algorithm are as follows:
**Step 1:** Initialize the model parameters based on the list provided in the table below (Table 1).
**Step 2:** Calculate ( $r_{off,i}$ ) based on the current clutch forces ($F_{c,i}$) and the sampled off-rate ($k_{off}$) from the long tailed distribution of ($\tau_{off}$), and find the clutch bound/engaged probability ($P_{b,i}$).
**Step 3:** Choose a random number for each clutch. If ($P_{b,i} > rand$), the clutch is considered engaged.
**Step 4:** Use the updated number of engaged clutches to solve for the substrate displacement ($x_s$) using the viscoelastic constitutive equation.
**Step 5:** Use force balance to simplify the equilibrium equations into the following summation equations for two motor clutch modules in each direction:

$$\sum_{i=1}^{n_c}\left(x_{c,i}^{+} + v_r^{+}\,dt - x_s\right)P_{b,i}^{+} = \sum_{i=1}^{n_c}\left(x_{c,i}^{+} + v_r^{+}\,dt - x_s\right)P_{b,i}^{+}$$

$$F_m\left(1 - \frac{v_r^{+} + v_r^{-}}{2v_m}\right) + F_r = k_c\sum_{i=1}^{n_c}\left(x_{c,i}^{+} + v_r^{+}\,dt - x_s\right)P_{b,i}^{+}$$

**Step 6:** Calculate the migration velocity and migration distance as:

$$v_m = \frac{v_r^{-} - v_r^{+}}{2}$$

$$d(t) = d(t-1) + v_m\,dt$$

$$F_r = k_m|\Delta x - \Delta y|$$

**Step 7:** Update the clutch forces($F_{c,i}$), myosin force ($F_m$) and membrane resistance force coupling the two dimensions ($F_r$) for the next simulation step ( $t + dt$ ).
**Step 8:** Sample a new $\left(\tau_{off}\right)$ from the power law distribution to calculate ($r_{off}$) in the next cycle.

| Variable | Parameter | Value | References |
|---|---|---|---|
| $\boldsymbol{k_a}$ | Additional stiffness | 0.1 – 10 pN/nm | Adjusted based on [2] |
| $\boldsymbol{k_l}$ | Long-term stiffness | 0.1 – 1 pN/nm | Adjusted based on [2] |
| $\boldsymbol{\eta}$ | Viscosity | 0.1-1000 pN-s/nm | This article |
| $\boldsymbol{\tau_s}$ | Substrate relaxation timescale | 1 s : Fast<br>1000 s : Slow | Fitting from experiments |
| $\boldsymbol{n_m}$ | Myosin motor number (on one module) | 200 | Adjusted based on [3, 4] |
| $\boldsymbol{n_c}$ | Clutch number (on one module) | 200 | Adjusted based on [3, 4] |
| $\boldsymbol{F_m}$ | Myosin force | 2 pN | Adjusted based on [3-5] |
| $\boldsymbol{F_b}$ | Characteristic clutch breakage force | 2 pN | Adjusted based on [3-5] |
| $\boldsymbol{r_{on}}$ | Association rate | 1 $s^{-1}$ | Adjusted based on [3-5] |
| $\boldsymbol{\tau_{min}}$ | Dissociation rate time constant distribution parameter | $1s^{-1}$ | This article |
| $\boldsymbol{\beta}$ | Glassy coefficient | 2.5 | This article |
| $\boldsymbol{k_c}$ | Clutch stiffness | 5 pN/nm | Adjusted based on [3-5] |
| $\boldsymbol{v_u}$ | Unloaded retrograde flow velocity | 120 nm/s | This article |
| $\boldsymbol{v_p}$ | Polymerization velocity | 120 nm/s | This article |

4. **Model predicts diffusive migration mode on wide range of substrate parameters:**
With an understanding of the hierarchy of timescales, we use our model to predict migration modes across a wide range of substrate parameters. Changing the substrate parameters ( $k_a$: $additional\ stiffness, k_l$: $long - term\ stiffness, \eta$: $viscosity$) can affect the previously discussed timescales $(\tau_s, \tau_l)$. Here, we use the glassy motor clutch model to obtain comprehensive phase diagrams of to show how the diffusivity exponent depends on the mechanical properties of the substrate. We choose $\beta$ = 1.5 to capture the glassy dynamics resulting from the long tail distribution of $\tau_{off}$, as increasing $\beta$ results in a fast-decaying tail, making it incapable of capturing differences in trap time and step size distribution tailed-ness with changing viscous properties. Consequently, the model cannot capture both sub- and super-diffusive migration modes, regardless of viscosity changes.
We find that viscosity affects cell migration in distinct ways depending on the elastic properties $(k_a, k_l)$, with the most significant impact at stiffnesses $(k_a \sim 1kPa)$. Stress relaxation is governed by $\sigma = k_l + k_a e^{-k_a t/\eta}$, for very small $k_a$ $(\sim 0.1\ kPa)$, and the time response is dampened due to the small coefficient multiplying the second term, which minimizes the effect of viscosity. In this case cells are unable to be trapped due to unstable clutches, leading to continuous migration steps and super-diffusive behavior. As $k_a$ increases, the effect of viscosity becomes more pronounced. For $k_a \cong 0.2\ kPa$, a small region in the phase diagram starts to show sub-diffusive migration. With further increase in $k_a \cong 1\ kPa$, corresponding to the stiffnesses of our experimental alginate substrates, we observe that higher viscosities result in sub-diffusive migration, while lower viscosities lead to super-diffusive migration. At this intermediate level of "additional" stiffness, the force transmission is optimal, allowing clutches to remain trapped for longer on slow-relaxing substrates. Conversely, on fast-relaxing substrates, the rapid decrease in overall stiffness destabilizes clutches, inhibiting trapping and resulting in longer migration steps characteristic of super-diffusive migration. As shown in SI Fig. 2, traversing the vertical line along increasing viscosity, the diffusivity decreases from $\alpha \cong 1.4$ to $\alpha \cong 0.6$, corresponding to the range observed in our experiments when cells were seeded on fast- vs. slow-relaxing substrates. For a further increase in $k_a$ (~10 kPa), viscosity effects diminish, and cells exhibit super-diffusivity for all parameter ranges due to rapid load and fail cycles that prevent trapping. Our previous experimental work also shows that viscosity only plays a role on soft substrates $(k_a \sim 1\ kPa)$ and cells on stiff substrates show no viscosity dependent effects [2]. Thus, sub-diffusion is only observed at higher viscosities on substrates with a stiffness of order of 1 kPa, whereas lower viscous properties and higher stiffness regimes predominantly results in super-diffusion. We next examine the role of

contractile forces in sustaining super-diffusive migration and how the impairment of contractility alters diffusivity.

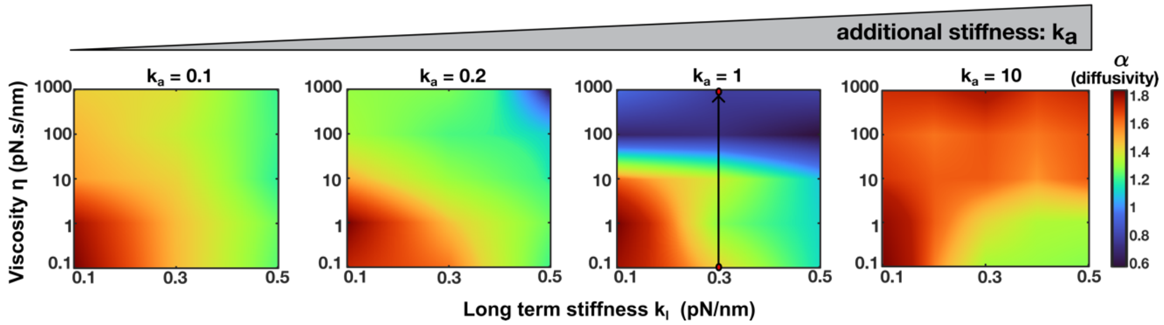


**Supplementary Figure 2:** Migration diffusivity on wide range of substrate parameters.

**6. Substrate relaxation and creep timescales:**

In the standard linear solid (SLS) model, which consists of a spring-dashpot-spring assembly, both the stress relaxation and creep responses offer insights into the viscoelastic properties of a material under different loading conditions. The stress relaxation timescale $\tau_{relax} = \frac{\eta}{k_a}$ captures how stress dissipates under a constant strain, while the creep timescale $\tau_{creep} = \frac{\eta(k_a+k_l)}{k_a.k_l}$ represents how the material progressively deforms over time when a constant load is applied. Here, η is the viscosity of the dashpot, and $k_l$ and $k_a$ are the long-term and additional stiffness respectively.

Despite these definitions, both timescales are closely related and generally fall within the same order of magnitude (SI Fig. 3). In real experiments, materials often exhibit both relaxation and creep behaviors simultaneously, meaning either timescale can provide a

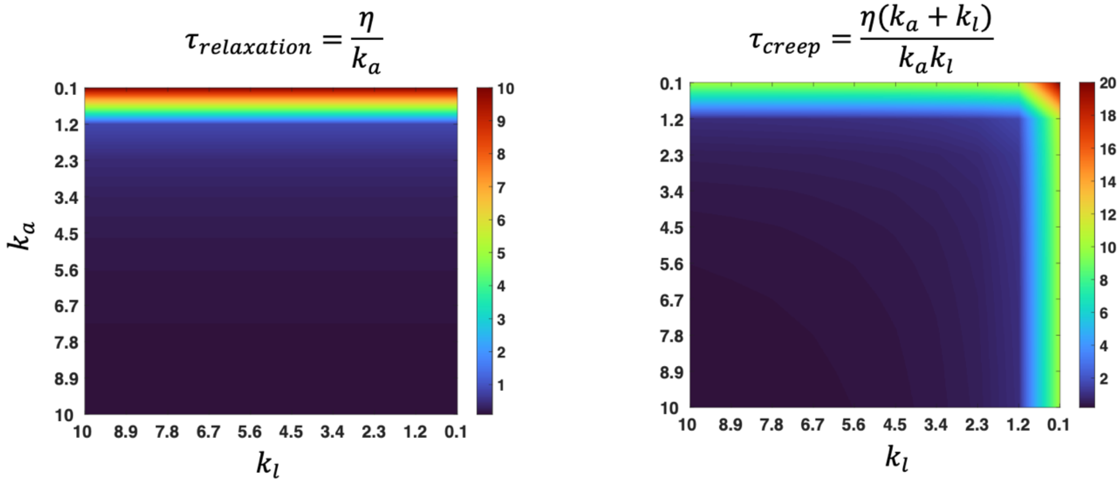


**Supplementary Figure 3:** Creep vs relaxation timescales as the substrate parameter changes.

valid approximation of the material's viscoelastic response. For simplicity in this work, we adopt the relaxation timescale, $\tau_s = \tau_{relax} = \frac{\eta}{k_a}$, given that rheology measurement on the alginate hydrogels measured stress relaxation timescale. This choice allows us to align our modeling with empirical observations, ensuring consistency with experimental conditions that focus on stress relaxation measurements.

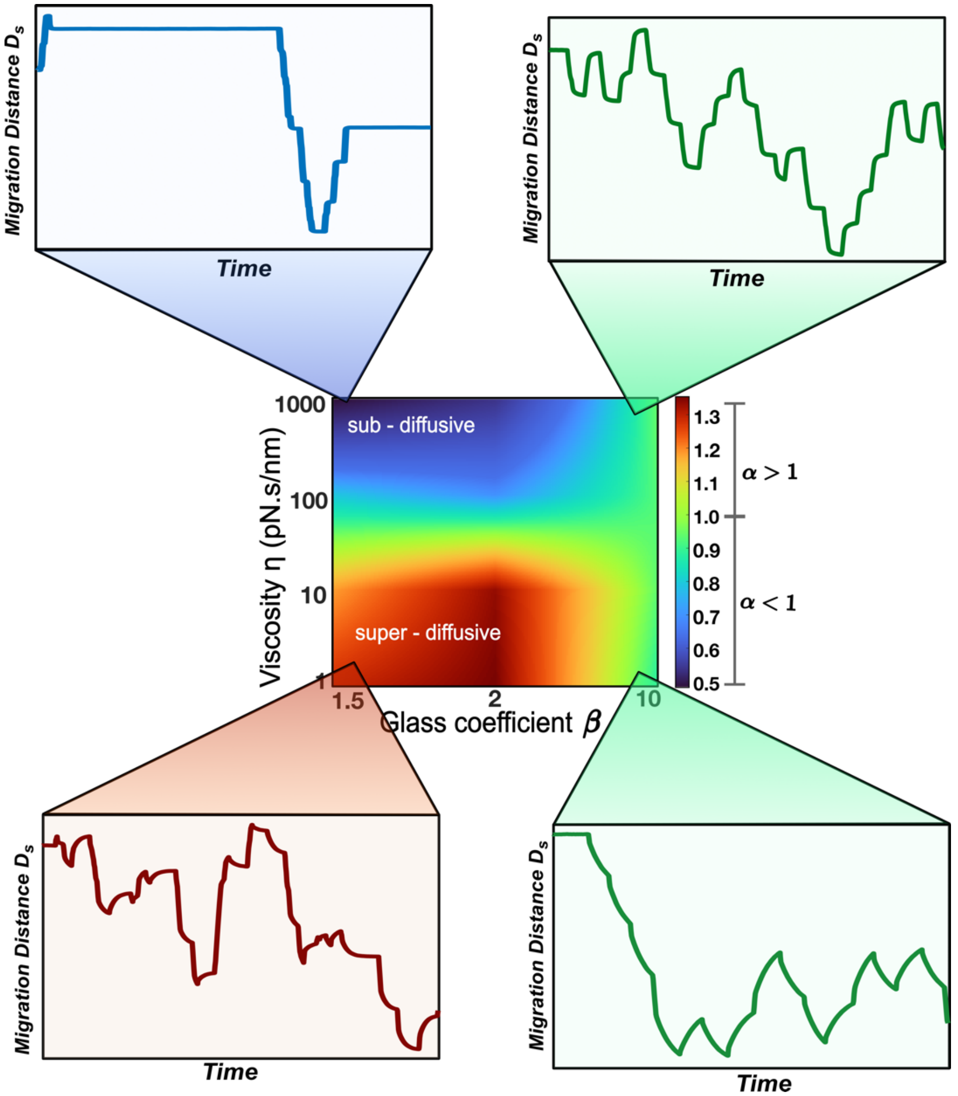


**Supplementary Figure 4:** Migration trajectories for glassy vs non-glassy model. Left represents migration trajectories for both fast (bottom) and slow (top) when glass coefficient value is small. Right side represents corresponding cases when glassiness is removed, and we see a periodic migration pattern for low and high viscosities.
